# Sex Differences in Oncogenic Mutational Processes

**DOI:** 10.1101/528968

**Authors:** Constance H. Li, Stephenie D. Prokopec, Ren X. Sun, Fouad Yousif, Nathaniel Schmitz, Paul C. Boutros, for the PCAWG Molecular Subtypes and Clinical Correlates Working Group, ICGC/TCGA Pan-Cancer Analysis of Whole Genomes Network

## Abstract

Sex differences have been observed in multiple facets of cancer epidemiology, treatment and biology, and in most cancers outside the sex organs. Efforts to link these clinical differences to specific molecular features have focused on somatic mutations within the coding regions of the genome. Here, we describe the first pan-cancer analysis of sex differences in whole genomes of 1,983 tumours of 28 subtypes from the ICGC Pan-Cancer Analysis of Whole Genomes project. We both confirm the results of exome studies, and also uncover previously undescribed sex differences. These include sex-biases in coding and non-coding cancer drivers, mutation prevalence and strikingly, in mutational signatures related to underlying mutational processes. These results underline the pervasiveness of molecular sex differences and strengthen the call for increased consideration of sex in cancer research.

Sex disparities in cancer epidemiology include an increased overall cancer risk in males corresponding with higher incidence in most tumor types, even after adjusting for known risk factors^1,2^. Cancer mortality is also higher in males, due in part to better survival for female patients in many cancer types, including those of the colon and head & neck^3^. Interestingly, female colorectal cancer patients respond better to surgery^4^ and adjuvant chemotherapy, though this is partially due to biases in tumour location and microsatellite instability^5^. Similarly, premenopausal female nasopharyngeal cancer patients have improved survival regardless of tumour stage, radiation or chemotherapy regimen^6^. There is a growing body of evidence for sex differences in cancer genomics^7-13^, but their molecular origins and clinical implications remain largely elusive.

Previous studies have mostly focused on protein coding regions, leaving the vast majority of the genome unexplored. We hypothesized that there are uncharacterized sex differences in the non-coding regions of the genome. Using whole genome sequencing data from the Pan-cancer Analysis of Whole Genomes (PCAWG) project^14^, we performed a survey of sex-biased mutations in 1,983 samples (1,213 male, 770 female) from 28 tumour subtypes, excluding those of the sex organs (**Supplementary Table 1**). We also excluded the X and Y chromosomes to focus on autosomal sex differences in cancers affecting both men and women, but there are known to be significant X-chromosome mutational differences between tumours arising in men and women^15^. Our analysis revealed sex differences in both genome-wide phenomena and in specific genes. These sex-biases occur not only at the pan-cancer level across all 1,983 samples, but also in individual tumour subtypes.

## Sex-biases in driver genes, mutation load and tumour evolution

We began by investigating sex differences in driver gene mutation frequencies, focusing on 165 coding and nine non-coding mutation events^16^ (**Supplementary Table 2**). We used proportions tests to identify candidate sex-biased events with a false discovery rate (FDR) threshold of 10%. These putative sex-biased events were then modeled using logistic regression (LGR) to adjust for tumour subtype, ancestry and age (**Online Methods**). We found several sex-biased pan-cancer driver events, including *CTNNB1* which was mutated in 5.0% more male-derived than female-derived tumours (male: 7.6%, female: 2.7%, 95% CI: 2.9-7.0%, prop-test q = 3.6×10^-4^, LGR q = 5.0 × 10^-3^; **Figure 1a, left**). *ALB* was also mutated in a larger proportion of male-derived tumours (male: 3.2%, female: 0.54%, 95% CI: 1.4-3.9%, prop-test q = 0.0038, LGR q = 6.5 × 10^-3^) while in contrast, *PTCH1* (male: 0.44%, female: 2.0%, 95% CI: 0.38-2.8%, prop-test q = 0.028, LGR q = 0.011) was mutated in more female-derived samples.

**Figure 1.**
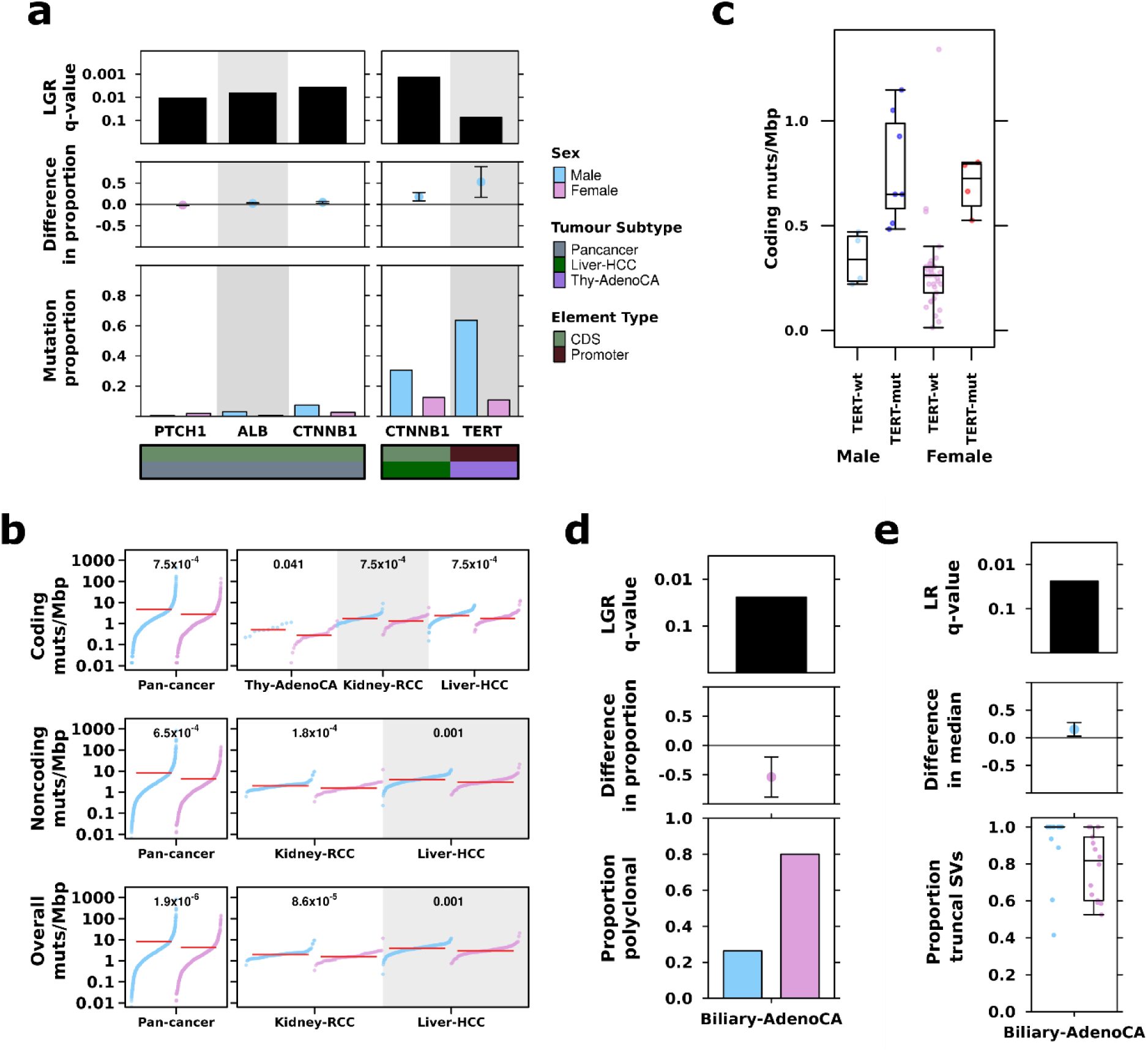
Sex biases in mutation frequency of driver genes, mutation prevalence and tumour evolution. **(a)** From top to bottom, each plot shows the logistic regression q-value for the sex effect; difference in proportion of mutated samples between the sexes, where blue denotes male-bias and pink denotes female-bias; and mutation proportion for each gene. Bottom covariate bars indicate mutation context and tumour subtype of interest. **(b)** The burden of somatic SNVs for coding, non-coding and overall mutation load. Linear regression q-values are shown. **(c)** Coding mutation load for thyroid adenocarcinoma samples compared by sex and presence or absence of TERT promoter mutations. **(d)** The proportion of polyclonal samples and **(e)** the proportion of truncal structural variants in biliary cancer.

We also identified tumour subtype-specific sex-biased driver mutations (**Figure 1a, right**). Similarly to the pan-cancer driver analysis, we first identified putative sex-biases using proportions tests and a 10% FDR threshold, and followed up with tumour subtype-specific logistic regression models (model descriptions in **Supplementary Table 1**). *CTNNB1* mutation frequency was sex-biased in liver hepatocellular cancer (Liver-HCC), again with more male-derived samples harbouring *CTNNB1* mutations: (male: 31%, female: 13%, 95% CI: 8.1-28%, prop-test q = 0.047, LGR q = 8.2×10^-3^, **Figure 1a, right**). This mirrors our previous finding of sex-biased *CTNNB1* mutation frequency in liver cancer from TCGA exome sequencing data, with similar effect sizes (male: 33% *vs.* female: 12%^11^). The largest sex-disparity was in a non-coding driver event in thyroid cancer (Thy-AdenoCA): *TERT* promoter mutations were observed in 64% of male-derived samples compared with only 11% of female-derived samples (95% CI: 18-89%, prop-test q = 6.9×10^-3^, LGR q = 0.074, **Figure 1a, right**), again supporting a previous finding^17^. Other putative sex-biased events were detected, but were not statistically significant after multivariate adjustment at present sample-sizes (**Supplementary Table 2**). These results demonstrate that mutation of key cancer-driving genes is sex-biased both within and across specific tumour subtypes.

Our previous work^12^ found sex biased mutation density across a number of tumour subtypes, including cancers of the liver, kidney and skin. We therefore investigated mutation density here to identify tumour subtypes where the cancer genomes of one sex accumulates more somatic single nucleotide variants (SNVs) than those of the other sex, and whether these sex-biases might be related to sex-biased driver gene mutation frequency. Returning to our statistical framework, we first used univariate tests to identify putative sex-biases, and then applied multivariate linear regression (LNR) on Box-cox transformed mutation load to adjust for possible confounders. The Box-cox transformation applies a power function to modify the shape of a variable’s distribution to better approximate a normal distribution (**Online Methods**). We also compared the total number of somatic SNVs and further divided mutations by coding and non-coding SNVs to determine whether sex-biases may be influenced by specific genomic contexts. Across all pan-cancer samples, we found higher mutation prevalence in male-derived samples in all three contexts (coding: difference in location = 0.41 mut/Mbp, 95% CI = 0.28-0.54 mut/Mbp, u-test q = 2.2×10^-10,^ LNR q = 7.5×10^-4^; non-coding: difference in location = 0.60 mut/Mbp, 95%CI = 0.43 - 0.80 mut/Mbp, u-test q = 7.9×10^-11^, LNR q = 6.5×10^-4^; overall: difference in location = 0.60 mut/Mbp, 95%CI = 0.42-0.79 mut/Mbp, u-test q = 7.5×10^-11^, LNR q= 1.9 × 10^-6^; **Supplementary Table 3**). These sex-biases remained significant even after adjusting for tumour subtype, ancestry and age in multivariate analysis (**Figure 1b, left**), demonstrating robust sex-biases in pan-cancer mutation prevalence across different contexts.

We also investigated somatic SNV burden in each of the 23 individual tumour subtypes with at least 15 samples, applying the same statistical approach with tumour subtype-specific models (model descriptions in **Supplementary Table 1).** We found sex-biased mutation load in three tumour subtypes (**Figure 1b, right**), with higher male coding mutation load in thyroid cancer (difference in location = 0.26 mut/Mbp, 95%CI = 0.12-0.43 mut/Mbp, u-test q = 0.028, LNR q = 0.041), and higher male load in hepatocellular cancer and kidney renal cell cancer (Kidney-RCC) in all three contexts (**Supplementary Table 3**). We compared the group rank differences of coding and non-coding mutation load between the sexes and found that in renal cell cancer, the differences were similar at 0.40 mut/Mbp for non-coding mutations and 0.37 mut/Mbp for coding mutations. In hepatocellular cancer however, the median sex-difference in non-coding mutation load was higher than the difference in coding mutation load (non-coding difference = 0.84 mut/Mbp *vs.* coding difference = 0.53 mut/Mbp). There is a similar effect in pan-cancer mutation load (non-coding difference = 0.60 mut/Mbp vs coding difference = 0.41 mut/Mbp) suggesting mutation context may play a role in sex-biased SNVs in some tumour subtypes.

To determine whether sex-biased mutation load may be associated with sex-biased driver gene mutation frequency, we focused on each driver gene and investigated SNV burden in the relevant tumour subtype. We did not find significant relationships between SNV burden and mutations in *PTCH1, ALB, CTNNB1* in pan-cancer analysis, nor was there an association for *CTNNB1* mutation in hepatocellular cancer. In thyroid cancer however, *TERT* promoter mutation was associated with increased coding mutation burden (median_*TERT-wt*_ = 0.26 mut/Mbp *vs* median_*TERT-mut*_ = 0.66 mut/Mbp, u-test p = 4.9×10^-6^). We used a linear regression model to determine if the sex-bias in coding mutation load could be explained by *TERT* mutation frequency and found this was indeed the case (linear regression p_TERT_ = 2.4×10^-5^, p_sex_ = 0.37, **Figure 1c**). In addition, we examined matched mutation timing data and found that of eleven samples with *TERT* promoter mutations, nine of these were truncal events, suggesting that an early sex-bias in *TERT* promoter mutation frequency is associated with sex-biased coding mutation load in this tumour subtype. Indeed, mutations in all sex-biased driver genes were overwhelmingly truncal events.

We then asked if these driver mutations might occur at different stages of tumour evolution between men and women, and started with tumour evolution structure. We compared the proportions of polyclonal *vs.* monoclonal tumours between the sexes and did not find significant sex differences in the proportions of polyclonal tumours bearing mutations in *PTCH1, ALB* or *CTNNB1* for sex-biased pan-cancer drivers, or in *TERT* promoter-mutated samples in thyroid cancer (**Supplementary Figure 1**). We did detect a putative bias in the proportion of polyclonal *CTNNB1*-mutated samples in hepatocellular cancer (80% of male-derived samples are polyclonal *vs*. 46% of female-derived samples, 95%CI = −0.019 – 0.70, prop-test p = 0.039), and accounted for polyclonality when comparing the timings of the mutations in these driver events. On subsequently examining the frequency of clonal *vs.* subclonal driver mutation events between the sexes, we found that while there were differences in the proportions of truncal mutations (*eg*: 100% of *TERT* promoter mutations were truncal events in male-derived *vs.* 50% truncal events in female-derived thyroid cancer patients), no comparisons were statistically significant.

Broadening beyond sex-biased driver mutations, we expanded our clonality analysis to perform a general survey of clonal structure and mutation timing across all tumour subtypes and mutations (**Supplementary Table 4)**. We found that female-derived biliary adenocarcinoma (Biliary-AdenoCA) tumours were frequently polyclonal, while most male-derived tumours were monoclonal (26% male-derived samples are polyclonal *vs*. 80% female-derived, 95% CI = 19 – 88%, prop-test q = 0.063, LGR q = 0.024; **Figure 1d**). In addition, we found intriguing evidence suggesting there may be sex-differences in the mutation timing of structural variants in this tumour subtype. Structural variants (SVs) in male-derived samples tended to be truncal events more frequently than in female-derived samples (median male percent truncal SVs = 100% *vs.* median female = 82%, u-test q = 0.081, LNR q = 0.024; **Figure 1e**). Though other comparisons did not reach our statistical significance threshold, we found some interesting trends that may merit future study, including in esophageal cancer (Eso-AdenoCA) where SVs in female-derived samples were more frequently truncal events while SVs in male-derived samples occurred more frequently in subclones (median male percent truncal SVs = 55%, median female = 100%; **Supplementary Figure 2**), and in medulloblastoma, where insertion-deletions (indels) were more frequently truncal events in female-derived samples than male (median male percent of truncal indels = 65%, median female proportion of truncal indels = 70%; **Supplementary Figure 3**). Our analysis of sex differences in tumour evolution identified some sex-biased events and also hint at putative sex-biases that should be further explored in future analyses.

## Sex-biases in genome instability and CNAs

Next, we examined percent genome altered (PGA), which provides a summary of copy number aberration (CNA) load. A proxy for genome instability, PGA is a complementary measure of mutation density to somatic SNV burden. While we did not find associations between sex and autosome-wide PGA, we observed sex-biases in the copy number burden for specific chromosomes (**Figure 2a**). In pan-cancer analysis, male-derived samples exhibited a slight but significant higher percent chromosome altered for chromosome 7 even after accounting for tumour subtype, ancestry and age using linear regression (median male PGA-7 = 5.4%, median female PGA-7 = 0.37%, difference in location = 0.0037%, 95%CI = 9.4×10^-4^–2.4×10^-3^%, u-test = 5.0×10^-3^, LNR q = 0.027; **Supplementary Table 5**). In individual tumour subtypes, we found sex-biased PGA in renal cell cancer (chromosomes 7 & 12) and hepatocellular cancer (chromosomes 1 & 16). By looking at copy number gains and losses separately, we additionally identified chromosomes with sex-biases in the burden of copy number gains and losses (**Supplementary Figure 4, Supplementary Table 5**), including sex-biased percent copy gained on chromosomes 5, 8 and 17 in pan-cancer samples.

**Figure 2.**
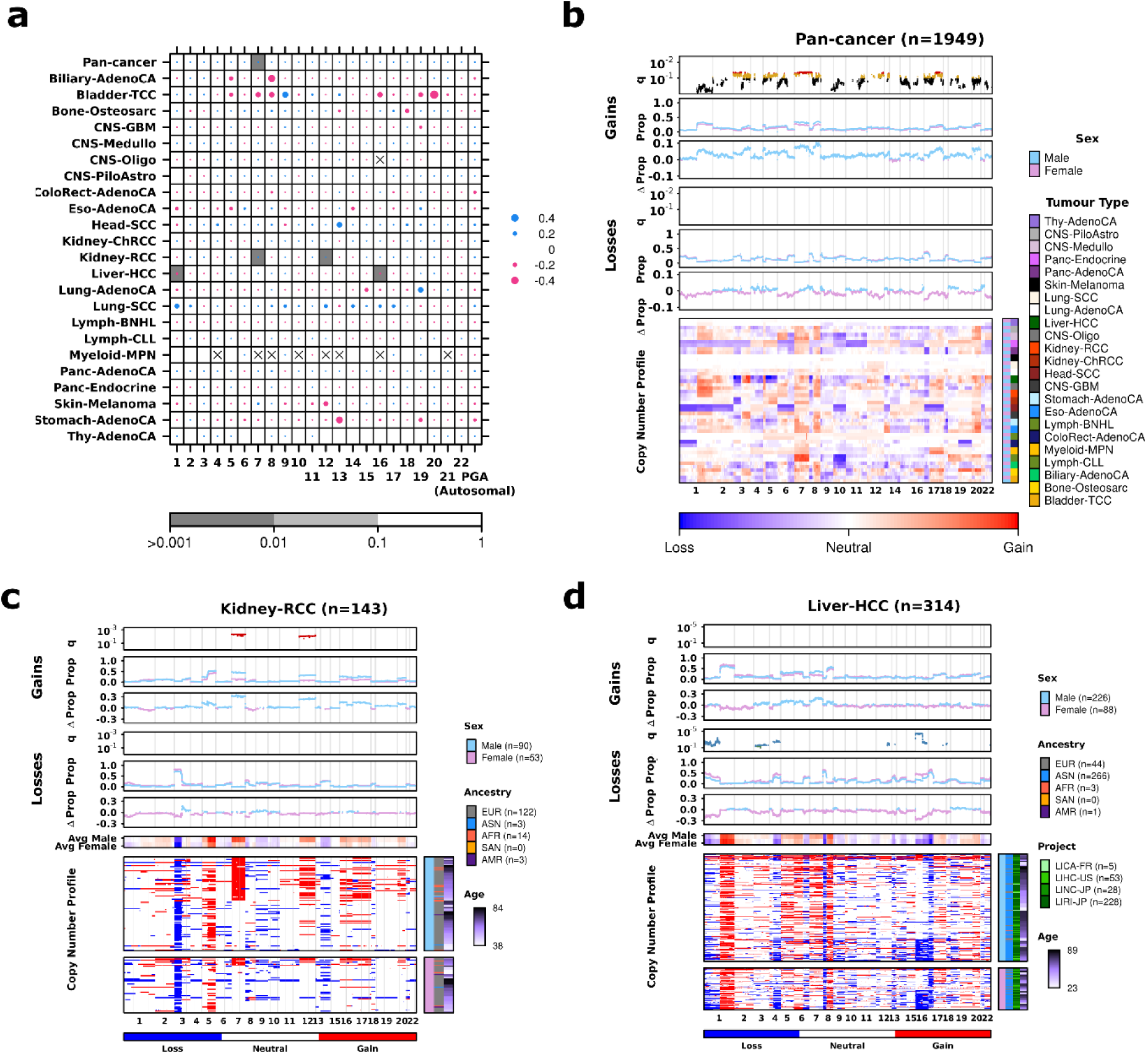
Sex-biases in percent chromosome altered are reflected in gene-specific events. **(a)** Dotmap showing association between sex and percent genome or chromosome altered, where dot size shows difference in median percent genome/chromosome altered between the sexes, and background shading shows q-values from linear regression. Sex differences in CNAs for **(b)** pan-cancer, **(c)** kidney renal cell cancer and **(d)** hepatocellular cancer. Each plot shows, from top to bottom: the q-value showing significance of sex from multivariate linear modeling with yellow (green) points corresponding to 0.1 < q < 0.05 and deep blue (red) points corresponding to q < 0.05; the proportion of samples with aberration; the difference in proportion between male and female groups for copy number gain events; the same repeated for copy number loss events; and the copy number aberration (CNA) profile heatmap. The columns represent genes ordered by chromosome. Light blue and pink points represent data for male- and female-derived samples respectively.

We next compared CNA frequency on the gene level to identify genes lost or gained at sex-biased rates. Across all pan-cancer samples, we found 4,285 sex-biased genes across 15 chromosomes (**Figure 2b, Supplementary Tables 6 & 7**, LGR q-value < 10%). These genes were all more frequently gained in male-derived samples than female with a difference in copy number gain frequency reaching ∼10% on chromosomes 7 and 8. Genes with male-dominated copy number gains include the oncogenes *MYC* (male gain frequency = 37% *vs.* female gain frequency = 28%, 95% CI = 5.2-14%, prop-test q = 2.5×10^-3^, LGR q = 0.068) and *ERBB2* (male gain frequency = 21% *vs.* female gain frequency = 16%, 4.7%, 95% CI = 1.1-8.3%, prop-test q = 0.041, LGR q = 0.088). The driver *CTNNB1* was also more frequently gained in male samples (male gain frequency = 8.9% *vs.* female gain frequency = 5.2%, 95% CI = 1.4-6.1%, prop-test q = 0.016, LGR q = 0.053), mirroring our finding of higher male pan-cancer mutation frequency on the SNV level for this oncogene. We did not find pan-cancer sex-biased copy number losses.

We repeated this analysis for every tumour subtype independently and found sex-biased CNAs in renal cell and hepatocellular cancer (**Supplementary Tables 6 & 7**). In renal cell cancer, the 1,986 sex-biased gains all occurred more frequently in male-derived samples, with differences in frequency up to 35% (**Figure 2c**). They spanned across chromosomes 7 and 12, agreeing with our finding of male-dominated genome instability in these chromosomes (**Figure 2a, Supplementary Figure 4**). In contrast to the male-dominated gain pan-cancer and renal cell findings, we found higher female frequency of copy number losses in hepatocellular cancer (**Figure 2d**). We identified 2,610 genes with higher copy number loss rates in female-derived samples. As observed in renal cell cancer, some of these losses span whole chromosomes, in this case chromosomes 3 and 16. Other sex-biased losses were found only across one chromosome arm (1p, 4q) or as focal events (*eg*. *PCDH9* on chromosome 13). The sex-biased gene-level events on chromosomes 1 and 16 agreed with the sex-biased genome instability findings but on returning to the PGA analysis, we found that chromosomes 3 and 4 had trending sex-biased genome instability (u-test q < 0.2, **Figure 1a, Supplementary Table 5**), suggesting that sex-biased PGA may guide identification of sex-biased CNAs on the gene level.

Thus, using sex-biased PGA as a guide, we more closely examined regions of interest in tumour subtypes of that did not have sex-biased CNAs in our general CNA analysis, but did have possible sex-biased genome instability (u-test q < 0.2): biliary cancer, B-cell non-Hodgkin lymphoma (Lymph-BNHL), chronic lymphocytic leukemia (Lymph-CLL) and melanoma (Skin-Melanoma). We found an additional 203 genes on the p-arm of chromosome 8 that were more frequently lost in female-derived samples in biliary cancer (**Supplementary Figure 5**). These copy number losses were 50% more common in female-derived samples and affect genes such as *DLC1*, a known tumour suppressor in hepatocellular cancer that is thought to play a similar role in gallbladder cancer^18^. While we did not identify additional sex-biased CNAs in non-Hodgkin lymphoma, chronic lymphocytic leukemia or melanoma, the sex-biased PGA results suggest these as regions of interest for future work. Thus in addition to sex-biased SNV events, we also identified sex-biased CNAs from this whole genome sequencing data.

## Sex biases in mutation signatures

We hypothesized that sex differences in mutation load and tumour evolution characteristics may be driven by varying mutational processes. In addition to single base substitution (SBS) signatures, which have been well annotated and linked to tumour aetiology^19,20^, we also examined doublet base substitution (DBS) and small insertion-deletion (ID) signatures. Sex differences in a mutational signature could shine insight on molecular differences between the sexes. For each of 47 validated PCAWG SBS, 11 DBS, and 17 ID signatures^21^, we performed a two-stage analysis. We first compared the proportions of signature-positive samples between the sexes; that is, we looked at the proportions of samples with any mutations attributed to the signature to determine whether there was a relationship between each signature and sex. Then, we focused on signature-positive samples and compared the percentage of mutations attributed to each signature between the sexes. For both analyses, we used univariate techniques to identify putative events and adjusted for additional variables using linear models.

At the pan-cancer level, we found eight signatures that occurred more frequently in one sex over the other (**Figure 3a, Supplementary Table 8**). In particular, SBS1 was more common in female-derived samples (89% of male-derived *vs.* 97% of female-derived, χ^2^-test q = 3.9×10^-10^, LGR q = 5.1×10^-7^) and was also associated with a higher percentage of mutations in these samples (male median percent mutations attributed to SBS1 = 8.4%, female median = 10%, u-test q =0.026, LNR q = 0.021). SBS1 is thought to be caused by deamination of 5-methylcytosine to thymine, resulting in base substitutions. Though it is correlated with age, our multivariate model accounts for this variable and the sex-bias remains even adjusting for age. SBS40 was also detected in a larger proportion of female-derived samples (42% of male-derived *vs.* 52% of female-derived, χ^2^-test q = 1.7×10^-4^, LNR q = 0.08), though we did not find a difference in the percentage of attributed mutations (u-test q = 0.17). Other sex-biased SBS signatures include SBS16, SBS17a and SBS17b, which were all more frequently detected in male-derived samples, and SBS40, which was more frequent in female-derived samples. These signatures are of unknown aetiology.

**Figure 3.**
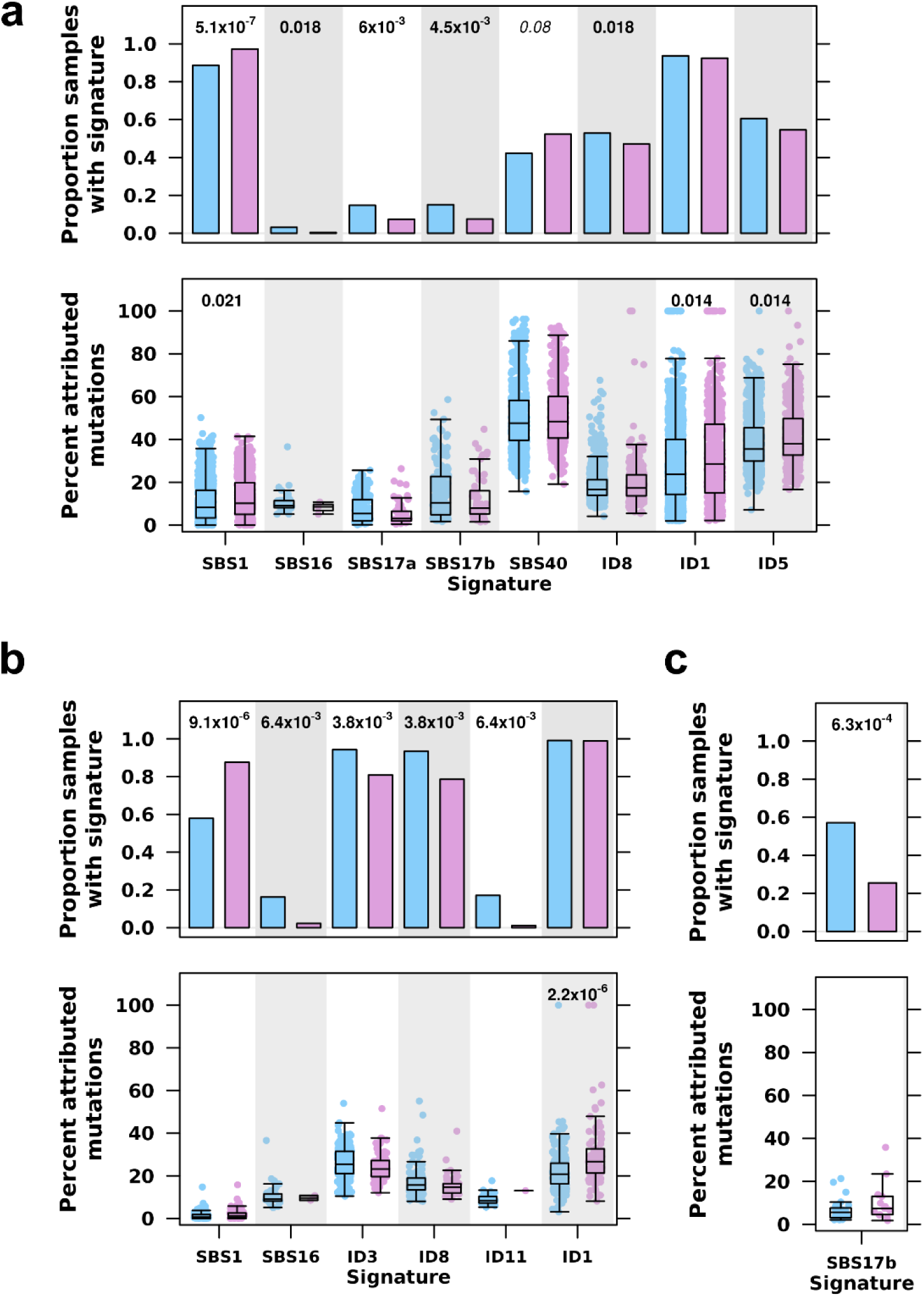
Sex differences in trinucleotide signatures related to mutational processes. Comparisons between proportion of signature positive samples shown in barcharts and proportion of attributed mutations shown in boxplots for **(a)** pan-cancer comparisons, **(b)** liver hepatocellular cancer, and **(c)** B-cell non-Hodgkin lymphoma. FDR-adjusted q-values for logistic regression (barplots) and linear regression (boxplots) shown only for significant comparisons. Blue shows male- and pink shows female-derived samples.

One ID signature was detected at different rates between the sexes, and two ID signatures had different rates of attributed mutations. ID8 occurred more frequently in male-derived samples (53% of male-derived *vs.* 47% of female-derived, χ^2^-test q = 0.068, LGR q = 0.018) though there was no difference in the percentage of mutations attributed to either signature. The aetiology underlying ID8 is not known, but this signature is thought to be associated with double strand break repair where ID8-asosciated mutations resemble those related to radiation-induced damage. Conversely, ID1 and ID5 were detected at similar frequencies between the sexes, but had higher percentages of attributed mutations in female-derived samples. Mutations associated with ID1 are thought to result from slippage during DNA replication and are associated with defective DNA mismatch repair, suggesting that while male- and female-derived tumours harbour defective DNA repair at similar rates, it is responsible for a larger proportion of mutations in female-derived tumours.

Since mutational processes are disease-specific, we repeated the mutational signatures analysis in each tumour subtype, again by first using univariate techniques to find putatively sex-biased signatures, and then using linear models to adjust for age and ancestry. We identified six sex-biased signatures in hepatocellular cancer (**Figure 3b, Supplementary Table 8**). Similar to our pan-cancer finding, we again detected female-dominated bias in the proportion of SBS1-positive samples (58% of male-derived *vs.* 88% of female-derived, χ^2^-test q = 3.5×10^-5^, LGR q = 9.2×10^-6^) male-dominated bias in and SBS16 (16% of male-derived *vs.* 2.2% of female-derived, χ^2^ q = 9.6×10^-3^, LGR q = 6.4×10^-3^). There were four sex-biased ID signatures in this tumour subtype: ID3 (94% of male-derived *vs.* 81% of female-derived, χ^2^-test q = 5.2×10^-3^, LGR q = 3.8×10^-3^), ID8 (93% of male-derived *vs.* 78.7% of female-derived, χ^2^-test q = 3.7×10^-3^, LGR q = 3.8×10^-3^) and ID11 (17% of male-derived *vs.* 1.1% of female-derived, χ^2^-test q = 3.7×10^-3^, LGR q = 6.4×10^-3^) occurred more frequently in male-derived samples. While ID1 was detected at similar rates between the sexes, a greater proportion of ID1-attributed mutations were found in female-derived than male-derived samples (male median percent mutations attributed to ID1 = 21%, female median = 27%, u-test q = 2.0×10^-5^, LR q = 2.2×10^-6^). As previously described, SBS1 and ID1 are associated with base deamination and defective DNA mismatch repair. ID3 is associated with tobacco smoke, and ID8 with double-stranded break repair. Taken together, sex-biases in the aetiology underlying the molecular landscape of hepatocellular cancer begin to emerge. In this tumour subtype, spontaneous or enzymatic deamination of 5-methylcytosine to thymine and defective mismatch repair occur more frequently in female patients and are also responsible for more mutations. Conversely, tobacco smoking is more common in male patients though the number of mutations attributed to tobacco smoke is not different between the sexes; this leads to more tobacco-associated male hepatocellular tumours.

In B-cell non-Hodgkin lymphoma, we identified a significant difference in the proportion of samples with SBS17b-attributed mutations (**Figure 3c, Supplementary Table 8**). More male-derived samples had mutations associated with this signature of unknown aetiology (57% of male-derived *vs.* 25% of female-derived, χ^2^-test q = 0.051, LGR q = 6.3×10^-4^). There were also several intriguing sex-differences in mutational signatures that did not meet our significance threshold. For instance, DBS2 accounts for a higher percentage of mutations in male-derived samples (male median percent mutations attributed to DBS2 = 50%, female median = 33%, **Supplementary Table 8**). DBS2’s association with tobacco smoking suggests that future insight in this signature may provide molecular explanations for the sex-specific associations between smoking and thyroid cancer risk^22^. As the aetiologies of these mutational signatures become better known, we can better approach the causes of molecular sex differences underlying cancer aetiology and progression. In particular, we may be able to discern environmental and lifestyle factors even in the absence of reported data, and connect known risk factors with newly described mutational processes.

Finally, to ensure that our findings were not skewed by differences in sequencing quality, we checked for sex-biases in quality control (QC) metrics. These included comparing the coverage, percentage of paired reads mapping to different chromosomes, and overall quality summary of both tumour and normal genomes. We mirrored our main analyses and used u-tests or χ^2^ tests and linear modeling to check each QC metric. We did not find sex-biases in any QC metric in pan-cancer or tumour subtype analysis after multiple adjustment except in raw somatic mutation calling (SMC) coverage. SMC coverage was higher in male-derived samples in six tumour subtypes including thyroid cancer and esophageal cancer, and was higher in female-derived samples in lung adenocarcinoma and B-cell non-Hodgkin lymphoma (**Supplementary Table 9**). While we do not find sex differences in comparing the SMC coverage pass/fail rates using a recommended minimum of 2.6 gigabases covered, it is prudent to consider sex-biased SMC in relation to our findings. There are also no sex-differences in the proportions of samples passing quality checks for any other QC metric (**Supplementary Table 9, Supplementary Figures 6**).

Our analysis of whole genome sequencing data from the PCAWG project uncovered sex differences in the largely unexplored non-coding autosomal genome. We found these biases in measures of mutational load, tumour evolution, mutational signatures, and at the gene level. While the majority of our findings describe pan-cancer differences, we have also uncovered an intriguing glimpse into tumour subtype-specific differences. These tumour subtype-specific results are limited by subtype sample size, and limited available annotation restricts the ability to account for confounding variables. It is important to consider these results in context of the multivariable models used, which do not directly capture characteristics such as tobacco smoking history or tumour stage at diagnosis. Future increases in sample size and robust associated annotation will allow for the detection of smaller effects and the control of more confounders. Nevertheless, our analyses of driver genes and copy number alterations suggest functional impacts of genomic sex-biases on the transcriptome and tumorigenesis. By using signatures to distinguish between mutations attributed to lifestyle factors such as smoking, we can better describe sex differences related to biological factors such as hormone activity. And despite low tumour subtype-specific sample numbers, our mutation timing and mutational signatures findings at both the pan-cancer and tumour-subtype level hint at underlying mutational processes that may give rise to molecular sex-biases. Combined with our previous work in whole exome sequencing, we present a landscape of sex-biases in cancer genomics and mutational processes (**Figure 4, Supplementary Figure 7**).

**Figure 4.**
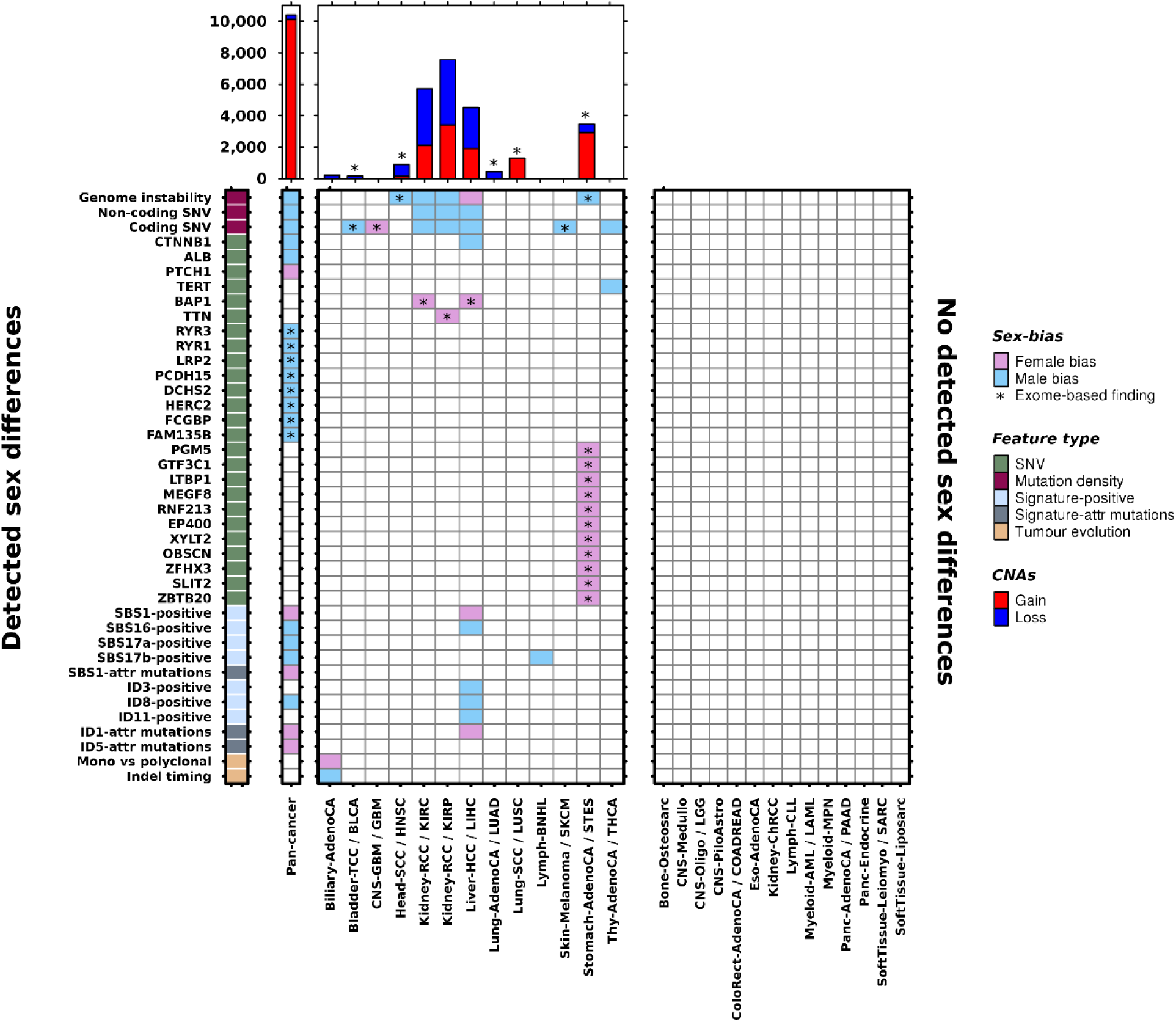
The landscape of sex differences in cancer genomics. Heatmap shows genomic features found to be sex-biased in pan-cancer analysis or in specific tumour subtypes. Results from both PCAWG and TCGA analyses are shown. Direction of sex-bias is shown in coloration denoting which sex has higher or more frequent aberration of the genomic feature. Top barplot shows union of genes found to be involved in sex-biased CNAs. Starred indicate findings exclusively from exome sequencing data (n=7,131) and unstarred indicate findings from PCAWG data (n=1,983).

It is becoming clear that sex differences occur across many mutation classes and the portrait of differences for each tumour subtype is a unique reflection of active mutational processes and tumour evolution. We have performed here the first pan-cancer analysis of sex differences in whole genome sequencing data and catalogued previously undescribed sex-biases. However, increased study of molecular sex differences in future large-scale sequencing efforts is needed to strengthen the findings we present here, to determine why men and women have molecularly different tumours, and to determine how this information can be leveraged to improve patient care.

## Acknowledgments

The authors thank all the members of the Boutros lab for insightful discussions. This study was conducted with the support of the Ontario Institute for Cancer Research to P.C.B. through funding provided by the Government of Ontario. This work was supported by the Discovery Frontiers: Advancing Big Data Science in Genomics Research program, which is jointly funded by the Natural Sciences and Engineering Research Council (NSERC) of Canada, the Canadian Institutes of Health Research (CIHR), Genome Canada and the Canada Foundation for Innovation (CFI). P.C.B. was supported by a Terry Fox Research Institute New Investigator Award and a CIHR New Investigator Award. This work was supported by an NSERC Discovery grant and by Canadian Institutes of Health Research, grant #SVB-145586, to PCB. The results described here are in part based upon data generated by the ICGC/TCGA Pan-Cancer Analysis of Whole Genomes Network: https://dcc.icgc.org/

## Author Contributions

CHL and PCB initiated the project. CHL, SDP, RXS, FY and NS analyzed data. PCB supervised research. CHL and PCB wrote the first draft of the manuscript, which all authors edited and approved. The PCAWG network provided variant calls and insightful commentary.

## Online Methods

### Data acquisition & Processing

Data was downloaded from the PCAWG consortium through Synapse. All data pre-processing was performed by the consortium as described^14^. Additional data-specific details are described below.

### General Statistical Framework

We followed a statistical approach as previously described in our previous work (Li et al). Briefly, for each genomic feature of interest, we used univariate tests first followed by false discovery rate (FDR) adjustment to identify putative sex-biases of interest (q < 0.1). Here, we use non-parametric univariate tests to minimize assumptions on the data. For putative sex-biases, we then follow up the univariate analysis with multivariate modeling to account for potential confounders using bespoke models for each tumour subtype. Model variables for each tumour context are described in **Supplementary Table 1** and were included based on availability of data (<15% missing), sufficient variability (at least two levels) and collinearity. Discrete data was modeled using logistic regression. Continuous data was first transformed using the Box-Cox family and modeled using linear regression. The Box-Cox family of transformations is a formalized method to select a power transformation to better approximate a normal-like distribution and stabilize variance. We used the Yeo-Johnson extension to the Box-Cox transformation that allows for zeros and negative values^23^:

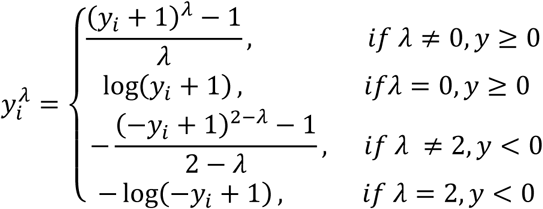

FDR adjustment was performed for p-values for the sex variable significance estimate and an FDR threshold of 10% was used to determine statistical significance. More detail is provided for each analysis below.

### Driver Event Analysis

We focused on driver events described by the PCAWG consortium^16^. Driver mutation data was binarized to indicate presence or absence of the driver event in each patient. Proportions of mutated genes were compared between the sexes using proportions tests for univariate analysis. A q-value threshold of 0.1 was used to select genes for further multivariate analysis using binary logistic regression. FDR correction was again applied and genes with significant pan-cancer sex terms were extracted from the models (q-value < 0.1). Driver event analysis was performed separately for pan-cancer analysis and for each tumour subtype.

### Clonal structure and mutation timing analysis

Subclonal structure and mutation timing calls were downloaded from Synapse (https://www.synapse.org/#!Synapse:syn8532425). Subclonal structure data was binarized from number of subclonal clusters per sample to monoclonal (one cluster) or polyclonal (more than one cluster). The proportion of polyclonal samples was calculated per sex and compared using proportion tests for both pan-cancer and tumour subtype analysis. The univariate p-values were FDR adjusted across all tumour subtypes to identify putatively sex-biased clonal structure. These cases were further scrutinized using logistic regression. A multivariate q-value threshold of 0.1 was used to determine statistically significant sex-biased clonal structure.

Mutation timing data classified SNVs, indels and SVs into clonal (truncal) or subclonal groups. The proportion of truncal variants was calculated for each mutation type (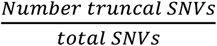, *etc*) to obtain proportions of truncal SNVs, indels and SVs for each sample. These proportions were compared between the sexes using two-sided Mann-Whitney *U*-tests and univariate p-values were FDR adjusted to identify putatively sex-biased mutation timing. Linear regression was used to adjust for confounding factors and a multivariate q-value threshold of 0.1 was used to determine statistically significant sex-biased mutation timing. The mutation timing analysis was performed separately for SNVs, indels and SVs.

### Mutation Load analysis

Consensus SNV calls were downloaded from Synapse (https://www.synapse.org/#!Synapse:syn7118450). Overall mutation prevalence per patient was calculated as the sum of SNVs across all genes on the autosomes and scaled to mutations/Mbp. Coding mutation prevalence only considers the coding regions of the genome, and noncoding prevalence only considers the noncoding regions. Mutation load was compared between the sexes using Mann-Whitney *U*-tests for both pan-cancer and tumour-type specific analysis. Comparisons with u-test q-values meeting an FDR threshold of 10% were further analyzed using linear regression to adjust for tumour subtype-specific variables. Mutation load analysis was performed separately for each mutation context, with pan-cancer and tumour subtype p-values adjusted together

### Chromosome and Genome Instability analysis

Consensus copy number data was obtained from Synapse (https://www.synapse.org/#!Synapse:syn8042880). Ploidy-adjusted calls were used to identify segments with copy number gains and losses. The number of bases in copy number gained or lost segments were summed per chromosome and divided by chromosome size to obtain percent chromosome gained and lost, respectively. All segments affected by a copy number aberration were also summed and treated in the same way to calculate percent chromosome altered. Percent copy number gained, lost, and altered were also calculated over the autosomes. These metrics were compared in pan-cancer and tumour-subtype analysis using u-tests to identify putatively sex-biased chromosome and genome instability, and putatively sex-biased events were further analysed using linear regression modeling. Genome instability analysis was performed separately for each tumour subtype with FDR adjustment performed over percent copy gained, loss and altered comparisons together.

### Genome-spanning CNA analysis

Consensus copy number data was processed to gain/neutral/loss calls per gene. The number of loss, neutral and gain calls were summed per sex, and assessed using univariate and multivariate techniques. For univariate analysis, proportional differences between the sexes for gains and losses were tested for each gene using proportions tests. After identifying candidate pan-cancer univariately significant genes, multivariate logistic regression was used to adjust ternary CNA data for sex, age, ancestry and tumour-type. The genome-spanning analysis was performed separately for losses and gains for each tumour subtype.

### Mutational Signatures analysis

The number of mutations attributed to each SBS, DBS and ID signature per sample was downloaded from Synapse (https://www.synapse.org/#!Synapse:syn8366024). For each signature, we compared the proportion of samples with any mutations attributed to the signatures (“signature-positive”) using *χ*^2^-square tests to identify univariately significant sex-biases. Signatures with putative sex-biases were further analysed using logistic regression.

We also compared the proportions of mutations attributed to each signature. The numbers of mutations per signature were divided by total number of mutations for each sample to obtain the proportion of mutations attributed to the signature. Mann-Whitney *U*-tests were used to compare these proportions. Putative sex-biased signatures were further analysed using linear regression after Box-cox adjustment.

Signatures that were not detected in a tumour subtype was omitted from analysis for that tumour subtype. Statistical analyses were performed for each set tumour subtype, but combining all SBS, DBS and ID signatures.

### Statistical Analysis & Data Visualization

All statistical analyses and data visualization were performed in the R statistical environment (v3.4.3) using the BPG^24^ (v5.9.8), car (v3.0-2) and mlogit (v0.2-4), packages.

