## SupplementaryFigures for "Sex Differences in Oncogenic Mutational Processes" (from another version of this preprint: version 1, 2019-01-23)

### Supplementary Figures 1-7

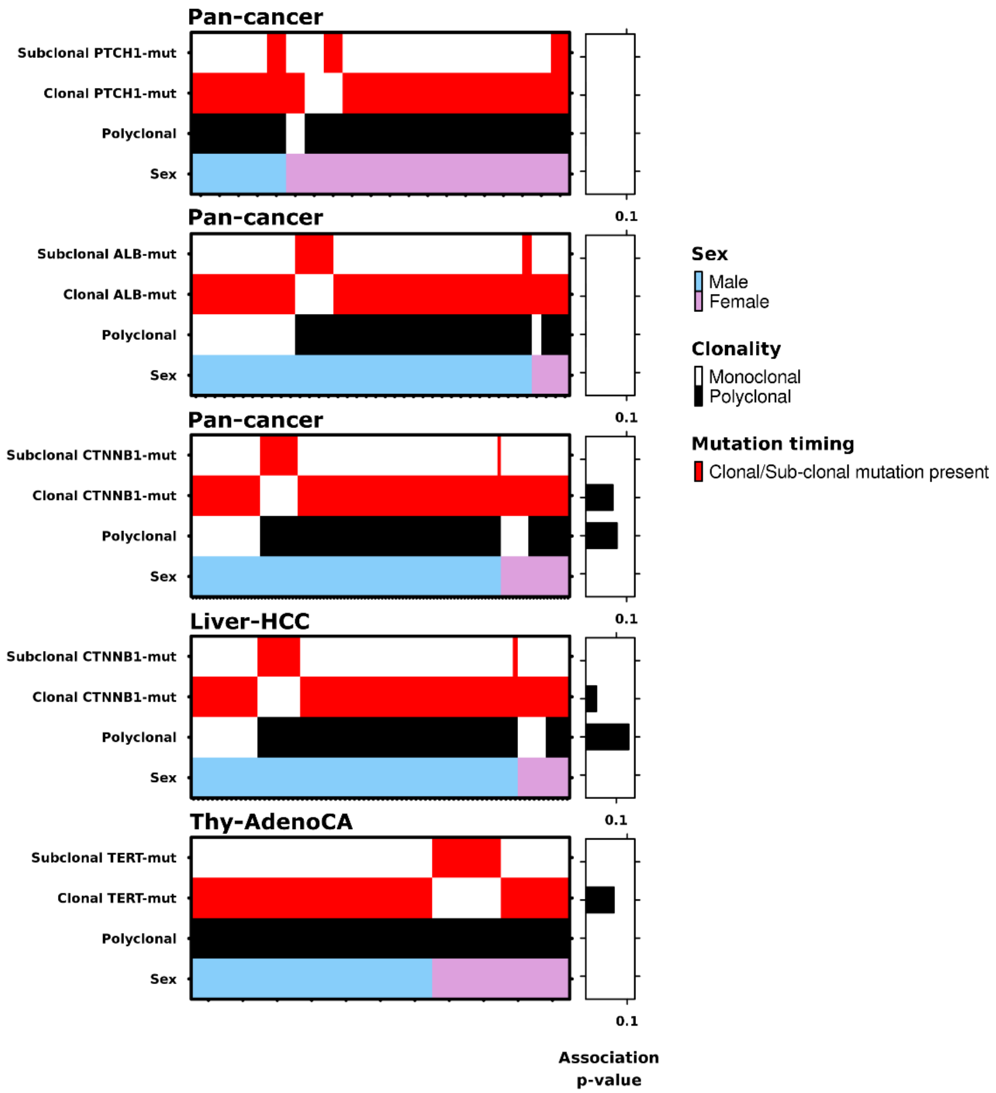


**Supplementary Figure 1. Mutation timing of sex-biased driver events.** For each driver event and tumour context, the heatmap shows the sex, polyclonality status, presence of clonal mutations, and presence of subclonal mutations for samples across the columns. Right barplots show p-values for the association between sex and the row of interest as tested by proportions tests.

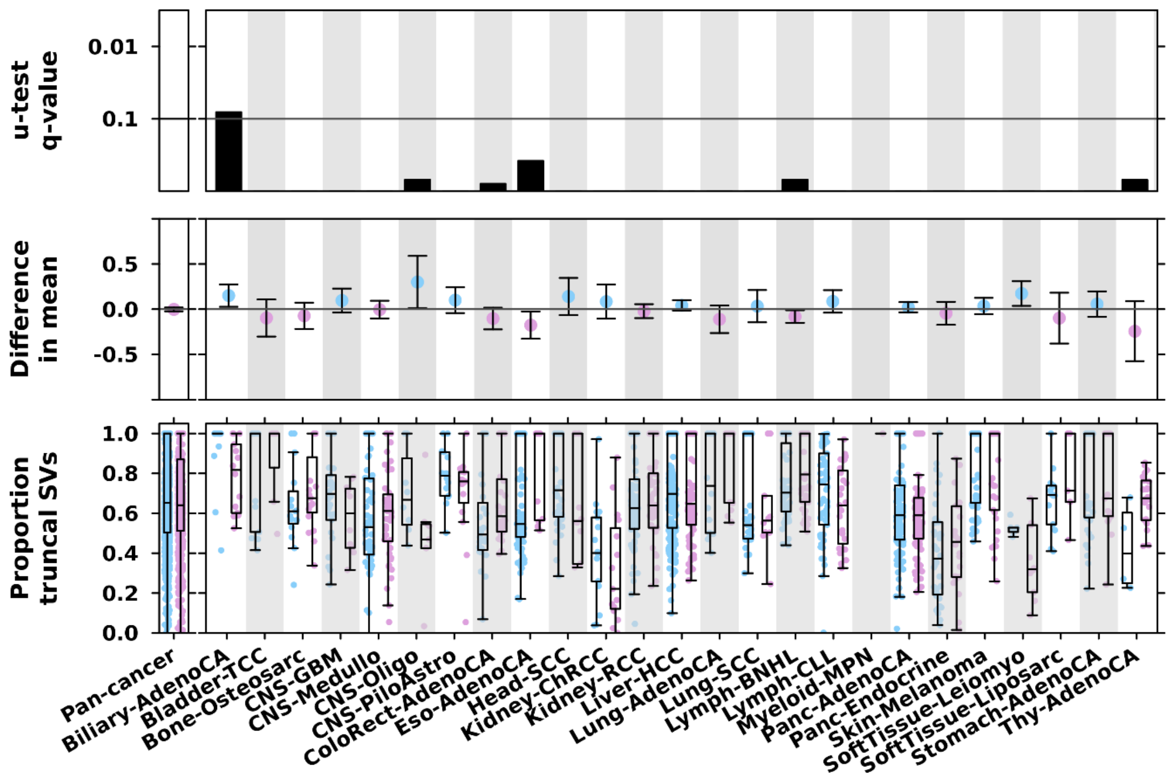


**Supplementary Figure 2. Structural variation mutation timing is not found to be sex-biased.** The proportion of truncal (*vs.* clonal) SVs is not statistically different between the sexes. From top to bottom: FDR-adjusted q-values from univariate *U*-tests, difference in means for proportion of truncal SNVs, and boxplots for proportion of truncal SNVs for each sample, where blue shows male- and pink shows female-derived samples.

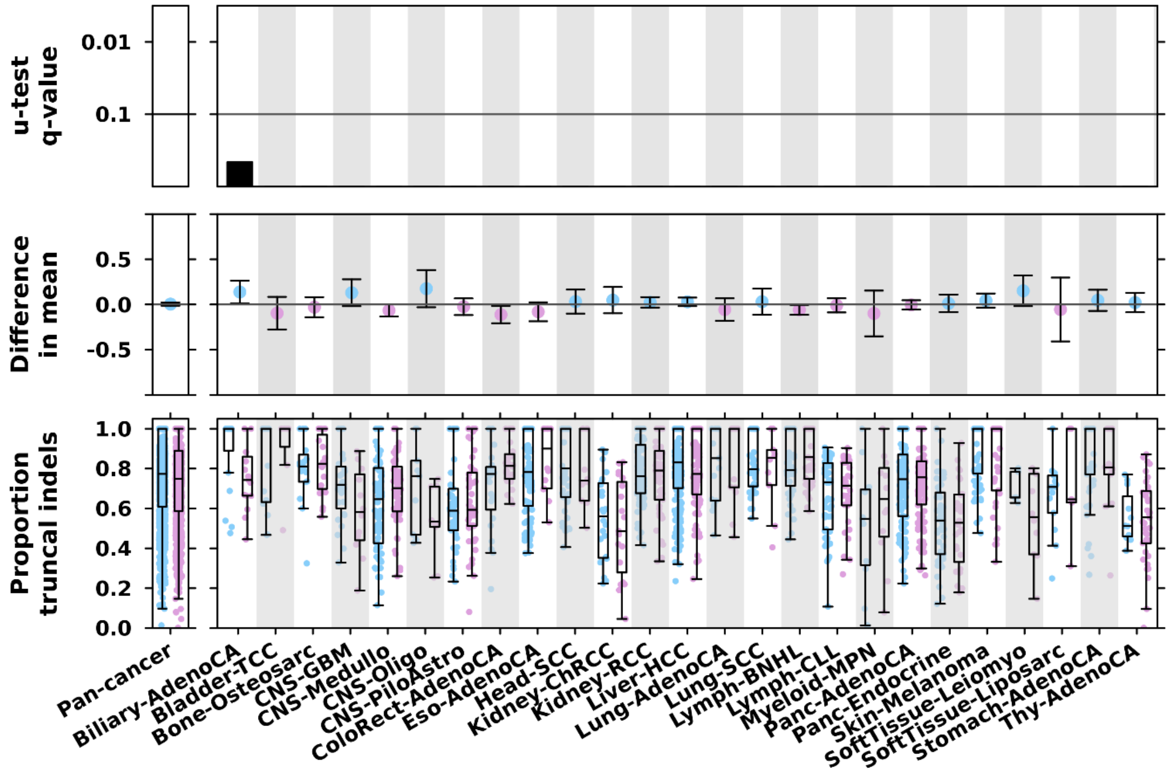


**Supplementary Figure 3. Indel mutation timing is putatively sex-biased in CNS-Medullo.** The proportion of truncal (*vs.* clonal) indels is sex-biased in CNS-Medullo. From top to bottom: FDR-adjusted q-values from univariate *U*-tests, difference in means for proportion of truncal SNVs, and boxplots for proportion of truncal SNVs for each sample, where blue shows male- and pink shows female-derived samples.

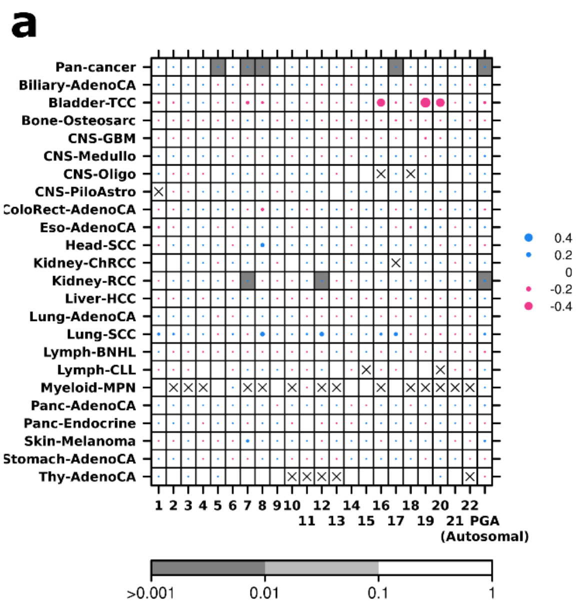


**Supplementary Figure 4. Sex-biased percent copy altered in each tumour subtype.** Dotmaps showing **(a)** percent genome or chromosome gained and **(b)** percent genome or chromosome lost. Dot size shows difference between median percent genome altered or percent chromosome altered. Background shading shows FDR-adjusted q-values from multivariate linear regression.

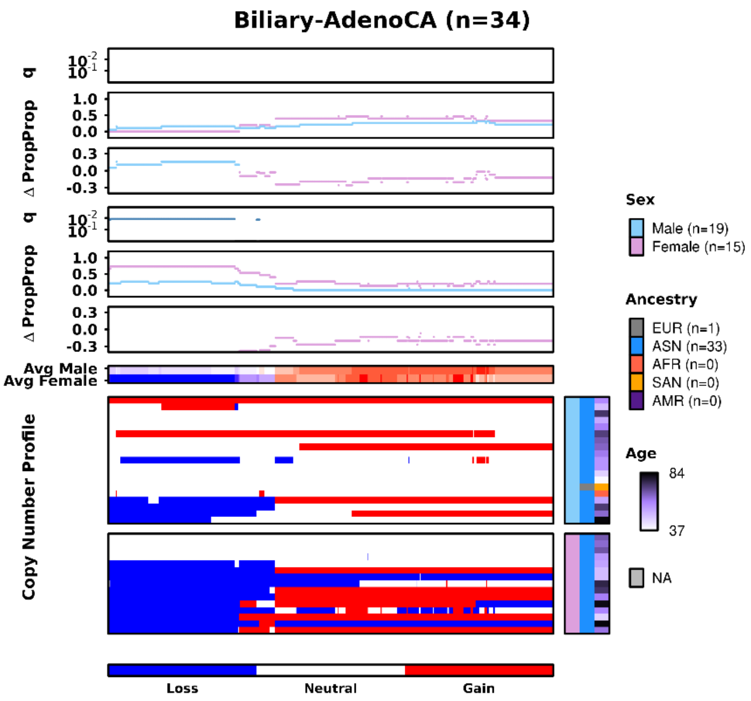


**Supplementary Figure 5. Sex-biased CNAs in Biliary Adenocarcinoma from PGA-guided analysis.** Each plot shows, from top to bottom: the q-value showing significance of sex from multivariate modeling on gene bins, proportion of samples with aberration, difference in proportion between male and female groups for amplifications; the same repeated for deletions and the CNA profile heatmap. The columns represent genes ordered by chromosome. The rows of the heatmap represent samples.

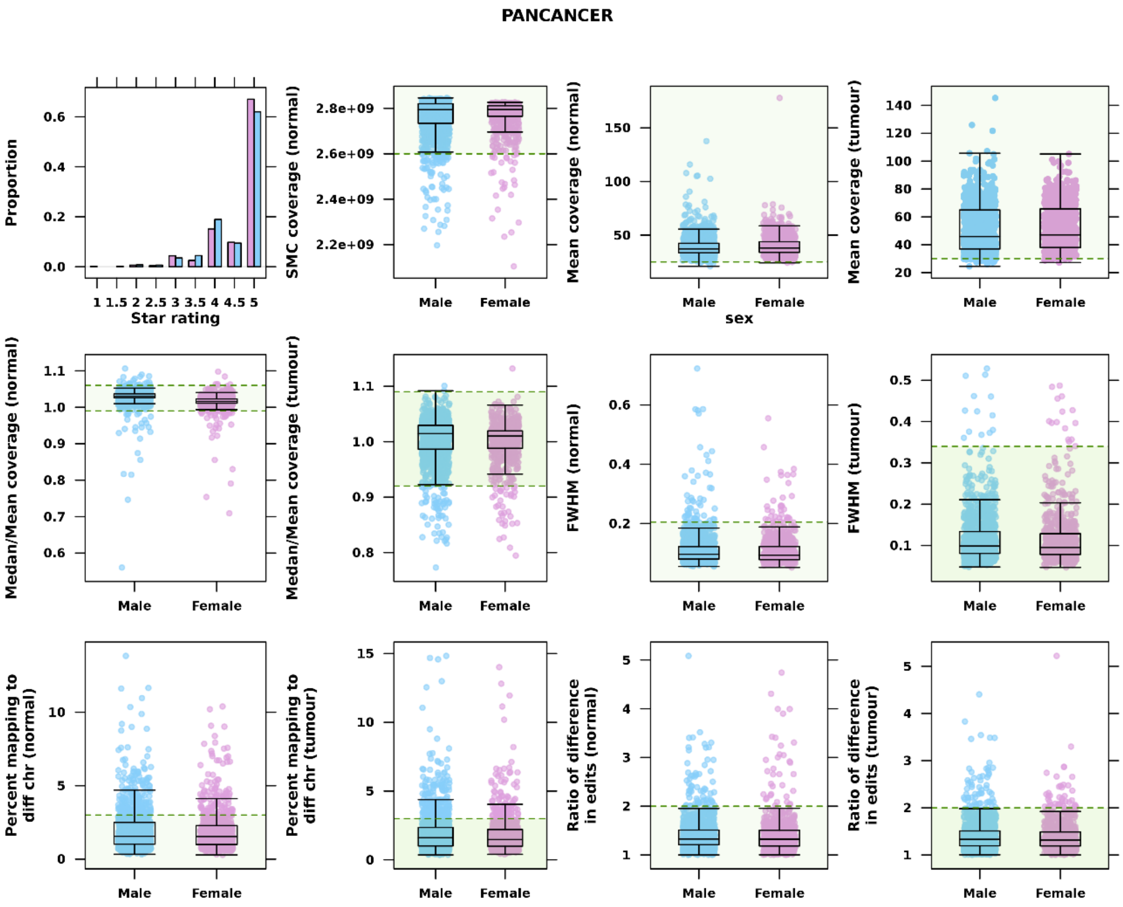


**Supplementary Figure 6. Quality control metrics by sex for pan-cancer samples.** Each plot shows the comparison between female- (pink) and male-derived (blue) samples for 12 quality control metrics. Green regions indicate PCAWG quality-pass thresholds.

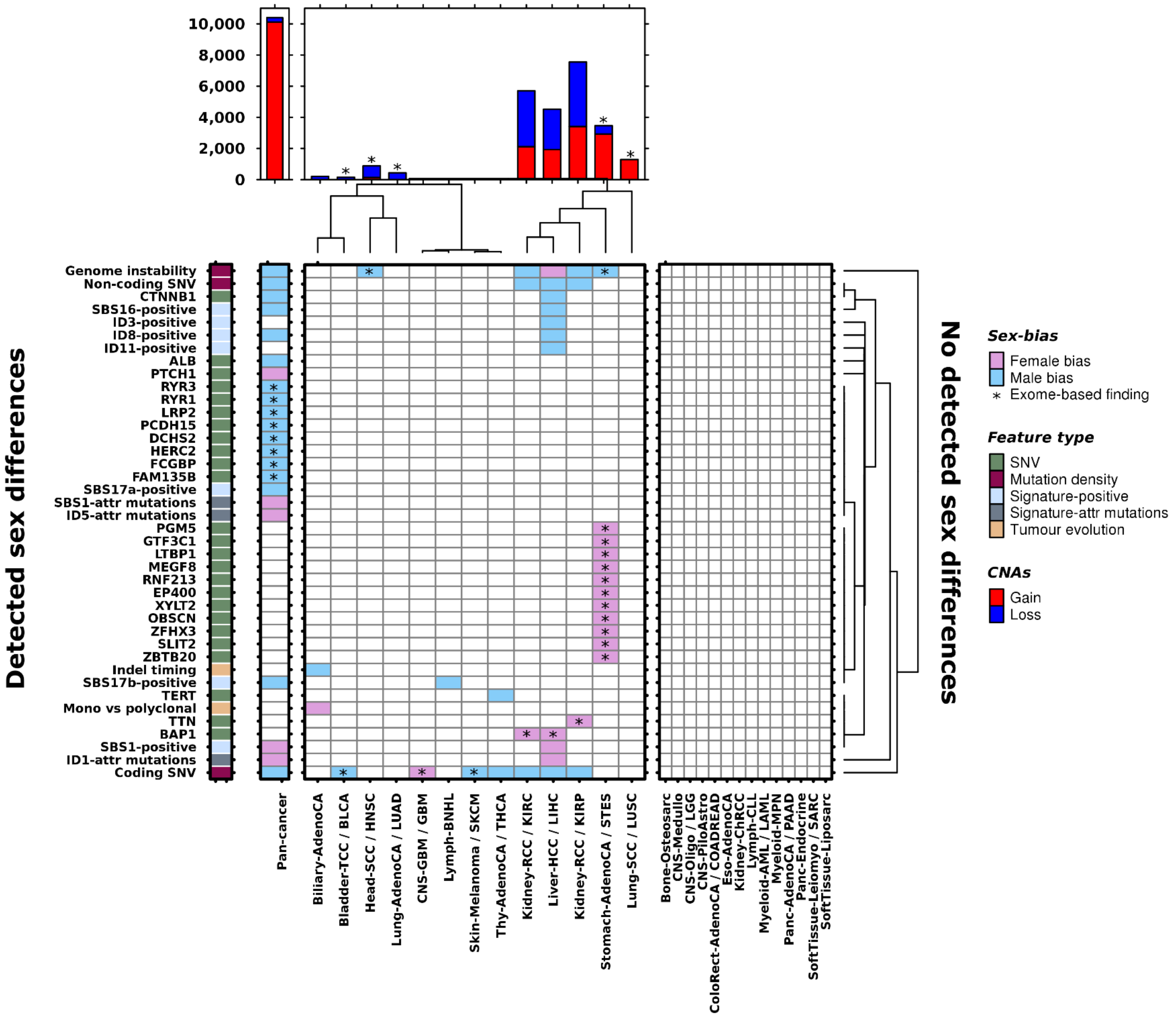


**Supplementary Figure 7. Clustered landscape of sex differences in cancer genomics.** Heatmap shows genomic features found to be sex-biased in pan-cancer analysis or in specific tumour subtypes. Results from both PCAWG and TCGA analyses are shown. Direction of sex-bias is shown in coloration denoting which sex has higher or more frequent aberration of the genomic feature. Top barplot shows union of genes found to be involved in sex-biased CNAs. Stars indicate findings exclusively from exome sequencing data (n=7,131) and unstarred indicate findings from PCAWG data (n=1,983). Diana clustering performed on rows and columns using Jaccard similarity coefficient.
